# Fecal proteomics of wild capuchins reveals impacts of season, diet, age, and, sex on gut physiology

**DOI:** 10.1101/2025.06.16.659980

**Authors:** Joseph D. Orkin, Alice Fournier, Daniel Young, Shasta E. Webb, Saul E. Cheves Hernandez, Katherine M. Jack, Fernando A. Campos, Antoine Dufour, Amanda D. Melin

## Abstract

Understanding how the physiology of free-ranging mammals is impacted by environmental stressors is a major focus of ecological research. However, the constraints of non-invasive sampling pose serious challenges to the acquisition of physiological data from most species of primates. As a result, little is known about how the gut responds to ecological stimuli at the cellular level in wild populations. Recent research has demonstrated that proteomics could fill this knowledge gap by sequencing and quantifying proteins directly from primate feces. In order to ascertain how the gut of free-ranging white-faced capuchin monkeys (*C. imitator*) is influenced by environmental heterogeneity, diet, age, and sex, we sequenced 45 fecal proteomes from 24 individuals from the Sector Santa Rosa population in Costa Rica, using liquid chromatography-tandem mass spectrometry with label-free quantification. Fecal proteins assigned to *C. imitator* were strongly localized to gut tissues and functionally enriched for digestive and immune functions. We identified 41 capuchin candidate proteins linked to seasonality, age, sex, and diet. We also quantified abundances of dietary fruit, dietary insects, helminth gut parasites, and gut microbes. Our results demonstrate the viability of using quantitative fecal proteomics in free-ranging populations of mammals to integrate host physiology, diet, and microbial ecology through non-invasive means.

## Introduction

Understanding how gut physiology and digestive function contribute to host health and plasticity is a driving question in primate ecology (Bergstrom et al., 2017; Campos et al., 2024; Lambert, 1998; Lambert & Rothman, 2015). Recent molecular advances have led to tremendous insights about these processes, particularly gut function (Rühlemann et al., 2024), health (Srivathsan et al., 2016), ecology (Orkin, Campos, et al., 2019), evolution (Amato et al., 2018), and ageing (Dasari et al., 2025). Despite all that is known, there remains a striking absence of quantitative data on how the gut functions at the cellular level in an ecological context. To a large extent, this results from the challenge of working with non-invasive biomaterials. Examining these questions in free-ranging primate populations–most of which are threatened with extinction–typically requires working with fecal samples (Arandjelovic & Vigilant, 2018; Orkin, Kuderna, et al., 2021). Luckily, feces provides a remarkable wealth of information. It is a heterogeneous mixture of tissues, including those from the host animal (e.g. gut epithelia, immune cells, and digestive enzymes), its recently consumed dietary items (e.g. fruit, leaves, and insects), gut microbiota, and parasitic/commensal organisms (Bădescu et al., 2017; Matthews et al., 2020; Moreno-Black, 1978; Orkin, Montague, et al., 2021; Tsutaya et al., 2021). Large scale metagenomic studies of free-ranging primate populations have used feces to explore these connections at the genetic level (Grieneisen et al., 2021; Rühlemann et al., 2024; Webb et al., 2023). However, genomes are not specific to particular cells and tissues, and DNA quantity does not reliably correspond to the amount of protein that has been translated in a given cell. As a result, it remains difficult to ascertain how free-ranging primates adapt to and are affected by fluctuating ecological circumstances at the cellular level.

Recently, the use of mass-spectrometry based proteomics to sequence proteins directly from primate feces has been highlighted as holding outstanding potential to quantify the abundances of gut proteins from both the host and exogenous sources (Tsutaya et al. 2021).

However, available data are from captive settings and efforts to integrate fecal proteomics into ecological research have been limited (Worsley et al., 2024). A major challenge is that, unlike DNA, peptide sequencing by mass spectrometry does not directly provide amino acid sequences. Instead, mass spectra are typically searched against known peptide sequence databases composed of proteome sequences from species expected to be found in the sample. The complexity of primate dietary and gut microbial ecology combined with the unevenness of available proteome assemblies makes it challenging to assemble a realistic metaproteomic database for free-ranging, non-model species in most contexts (Tsutaya et al., 2021).

Nonetheless, because proteomes are tissue specific, in the appropriate study system direct measurements of protein abundances from the cells present in feces–both from host and exogenous sources–could be obtained by quantifying peptide spectral intensities. As such, fecal proteomics could be used to identify context specific differential abundances of proteins relevant to primate health, ageing, dietary ecology, physiology, metabolism, and behavior. Proteins also offer other advantages in the context of molecular ecology. They are stable and resistant to degradation, allowing for them to be sequenced even from the poorly preserved and desiccated samples common to field conditions (Tsutaya et al., 2021), and unlike with DNA, the risk of contamination is minimal because peptide sequences assigned to foreign organisms or unexpected tissues (like human skin cells) can simply be excluded from downstream analysis.

Ultimately, leveraging proteomics to examine how the gut physiology of free ranging primates interacts with their behavioral ecology, health, and ageing, requires working with a population that has substantial pre-existing genomic, metagenomic, and ecological data.

One such population is the white-faced capuchin monkeys (*Cebus imitator*) of Sector Santa Rosa (SSR) in northwestern Costa Rica. Decades of behavioral and molecular research at the site have resulted in detailed records of plant phenology along with the dietary and gut microbial compositions from dozens of well-known, habituated individuals (Fedigan & Rose-Wiles, 1996; Melin et al., 2020; Orkin, Webb, et al., 2019; Webb et al., 2023). After more than 40 years of intensive study (Fragaszy et al., 2004; Melin et al., 2020), it is evident that capuchin survival depends on adapting to and overcoming the stresses of intense seasonal resource scarcity in this challenging environment (Campos & Fedigan, 2009; Orkin, Montague, et al., 2021). SSR occurs within a tropical dry forest biome, with strong seasonality in rainfall and temperature (Janzen, 2002). There is a marked dry season between November and May during which precipitation is almost entirely absent; in contrast, heavy amounts of rain (observed up to 3498 mm) can fall during the wet season, which extends throughout the remainder of the year (Campos, 2018; Campos & Fedigan, 2009). Fruit resources are also seasonally abundant at SSR, typically punctuated by a decline in fruit biomass during June and July (Orkin, Campos, et al., 2019). The selective effect of these conditions on resident primates can be profound. Periods of intense drought, often exacerbated by el niño conditions, have led to increases in infant mortality (Campos et al., 2020) and diminished fruit resources have been associated with loss of muscle mass (Bergstrom et al., 2017) and compensatory insect consumption (Orkin, Campos, et al., 2019). In response to these challenges, capuchins at SSR have undergone selection at loci related to seasonal resource scarcity, including genes involved in sugar metabolism, water balance, kidney function, and muscular wasting (Orkin, Montague, et al., 2021). Substantial shifts in gut microbial diversity–particularly among the dominant taxa *Bifidobacteria* and *Streptococcus*–have been observed in response to seasonality and variable levels of fruit and insect composition (Orkin, Campos, et al., 2019). Fluctuations in the gut diversity of parasitic helminths have been observed across seasons as well (Pinto et al., 2023) adding a further signal of season stress. Furthermore, capuchins live exceptionally long lives–reaching 54 years in captivity (Melin et al., 2020)–and the SSR population is widely studied in the context of social and ecological ageing processes (Campos et al., 2024; Sadoughi et al., 2025). Given the context of seasonal stress and muscular wasting, age-associated patterns of increasing frailty and inflammation throughout life are of particular concern in this population.

Despite all that is known about the ecology, genetics, and microbiomes of free-ranging primates like the capuchins at SSR, it has not yet been possible to integrate these data with the cellular functioning of the gut in a quantitative manner. We leveraged fecal proteomics to examine how the white-faced capuchin gut physiology responds to and is affected by fluctuating ecological circumstances. Specifically, we sequenced whole proteomes of 45 fecal samples from 24 individual white-faced capuchins from SSR, using liquid chromatography-tandem mass spectrometry (LC-MS/MS) with label-free quantification (LFQ). We use these data to: 1) classify the cellular composition and functional enrichment of the *C. imitator* gut proteome; 2) quantify the abundance of non-host proteins present in the *C. imitator* gut, including those from dietary items, gut microbiota, and intestinal parasites; and 3) demonstrate how the abundances of different proteins in the *C. imitator* digestive tract are affected by ecological and host-specific variables. To our knowledge, this is the first study to use quantitative fecal proteomics on a free-ranging population of mammals. We reveal a path forward toward the integration of host physiology and dietary and microbial ecology through non-invasive means.

## Methods

### Study population and fecal sampling

White-faced capuchin monkeys are medium-bodied, arboreal primates that consume an omnivorous diet dominated by fruit and invertebrate foods foraged from all forest strata (Williamson et al., 2021). While they inhabit a geographic distribution ranging from Honduras to Panama, sampling was conducted solely from the SSR population, which is located within the Área de Conservación Guanacaste in Costa Rica (10.84, -85.62). These individuals are known to long-term behavioral observers by distinct visual features, allowing for confident replicate sampling. Since 2014, fresh fecal samples from the SSR capuchin population have been collected at regular intervals and biobanked at -80°C at The University of Calgary (Webb et al., 2023). Researchers wearing gloves collected samples from the forest floor immediately following defecation and placed them into sterile 2 ml tubes, which were then stored on ice packs until being placed in liquid nitrogen at the field station before shipment. Fecal samples were visually inspected for the presence of dietary fruits and insects known to researchers. We selected paired fecal samples from 24 individuals collected once during the end of the dry season (April 29 - May 7, 2014) and once during the peak of wet season (July 13 - August 5, 2015), in order to represent the ecological contrast at SSR.

### Protein extraction and preparation

100 µg of feces was placed in a 1% SDS / 0.5M EDTA buffer and underwent 20 minutes of bead beating at 30 hz to homogenize and lyse the sample. Samples were centrifuged at maximum speed for one minute and then sonicated twice on ice for 5-10 seconds with a probe sonicator. Following ten minutes of 4°C centrifugation at 14,000 x g to remove cellular debris, supernatants were collected. Total protein quantification was done with a Pierce BCA Protein Assay Kit (Thermo Scientific) measured against standard curves using the microplate procedure. Proteins were purified and concentrated using trichloric acid (TCA) precipitation and filter-aided sample preparation (FASP). 10 mM of DTT was added to 200 µg of protein from each sample and incubated at 37°C for thirty minutes. Reduced proteins were transferred to a Micron YM-30 filter, washed with 200 µL of Urea-Tris and incubated with 100 µL of iodoacetamide. Samples were washed again with Urea-Tris and twice with 100 µL of 50 mM ammonium bicarbonate. Peptides were digested overnight at 37°C with 20 µg of trypsin (Promega) and 80 µL of 50 mM ammonium bicarbonate. Recovered peptides were washed as above and eluted with 40 µL of 0.5 M NaCl, then lyophilized and resuspended in 1% Formic acid. C-18 solid-phase extraction (CPE) cleanup was conducted to remove any final contaminants and load samples for LC-MS/MS.

### Liquid Chromatography

All samples were analyzed at the Southern Alberta Mass Spectrometry (SAMS) core facility. Tryptic peptides were analyzed on an Orbitrap Fusion Lumos Tribrid mass spectrometer (Thermo Fisher Scientific) operated with Xcalibur (version 4.0.21.10) and coupled to a Thermo Scientific Easy-nLC (nanoflow Liquid Chromatography) 1200 system. 2 μg of Tryptic peptides were loaded onto a C18 trap (75 μm × 2 cm; Acclaim PepMap 100, P/N 164946; Thermo Fisher Scientific) at a flow rate of 2 μl/min of solvent A (0.1% FA and 3% acetonitrile in LC–MS grade water). Peptides were eluted using a 90 gradient from 5 to 40% (5% to 28% in 105 min followed by an increase to 40% B in 15 min) of solvent B (0.1% FA in 80% LC–MS grade acetonitrile) at a flow rate of 0.3 μL/min and separated on a C18 analytical column (75 μm × 50 cm; PepMap RSLC C18; P/N ES803; Thermo Fisher Scientific). The mass spectrometer (Orbitrap Lumos) was calibrated before each batch and 100 fmol of Pierce BSA protein digest (PI88341) was injected to control the performance of the liquid chromatography and the mass spectrometer before the samples were acquired.

### Tandem Mass Spectrometry

We acquired the data using a label free quantification (LFQ) approach. Peptides were electrosprayed using 2.1 kV voltage into the ion transfer tube (300 °C) of the Orbitrap Lumos operating in positive mode. The Orbitrap performed a full MS scan at a resolution of 120,000 FWHM to detect the precursor ion having m/z between 375 and 1,575 and a + 2 to + 7 charge. The Orbitrap automatic gain control (AGC) and the maximum injection time were set at 4 ×105 and 50 ms, respectively. The Orbitrap was operated using the top speed mode with a 3 second cycle time for precursor selection. The most intense precursor ions presenting a peptidic isotopic profile and having an intensity threshold of at least 5,000 were isolated using the quadrupole and fragmented by higher-energy collisional dissociation (HCD, 30% collision energy) in the ion routing multipole. The fragment ions (MS2) were analysed in the ion trap at a rapid scan rate. AGC and the maximum injection time were set at 1 X 104 and 35 ms, respectively, for the ion trap. Dynamic exclusion was enabled for 45 s to avoid the acquisition of the same precursor ion having a similar m/z (plus or minus 10 ppm).

### Protein Database Searches and Quantification

We constructed a protein database using both the *Cebus imitator* proteome and 71 other species of microbes, invertebrates, and plants (Table S1) based upon the known dietary ecology and gut microbiota of free-ranging *Cebus imitator* at SSR. While there are no UniProt reference proteomes available for the specific invertebrates consumed by *C. imitator*, we selected those of six closest relatives in UniProt (*Aristophania vespae*, *Heliothis virescens*, *Manduca sexta*, *Nezara viridula*, *Trachymyrmex cornetzi*, and *Vespula vulgaris*). Given the paucity of proteomes available from fruiting plants, we selected only one proteome, *Ficus carica*. Furthermore, given the large size of plant genomes, including additional plant proteomes would have been computationally limiting. Additionally, we included the proteomes from available helminth and protozoan species within genera known to parasitize *C. imitator* at SSR: *Strongyloides venezuelensis* and *Giardia intestinalis*.

Raw spectra files were searched against the database using the Data Dependent Analysis (DDA) with the label-free quantification pipeline in FragPipe (v22.0) (Kong et al., 2017). A set of decoy proteins, prefixed with rev_ was added to the database along with the set of likely laboratory contaminant proteins from the crapOME repository. For MSFragger (v4.1) searches, non-default settings included: precursor mass tolerance of +/- 10 PPM, minimum peptide length of 6, maximum peptide mass range of 6,600, up to 3 variable modifications allowed per peptide, including oxidation of methionine, N-terminal acetylation, and deamidation. Enzymatic cleavage with “stricttrypsin” allowed up to 2 missed cleavages. Validation was run with Crystal-C (Chang et al., 2020) and PeptideProphet (Ma et al., 2012), and FDR filters (protein, peptide, psm, and ion) were set to 0.05. Posttranslational modifications were identified with PTM-Shepherd (Geiszler et al., 2021). MS1 quantification was performed with IonQuant. Intensities were normalized across runs and MaxLFQ min ions were set to 2, and the top 3 ions were used to quantify intensity.

### Data normalization and cleaning

Peptide intensities were normalized in MSstats (v4.6.5) (Kohler et al., 2023) by log transformation and equalizing median values. Protein groups identified as contaminants and keratins (from potential human skin cells) were removed from the dataset. In order to improve the annotation of proteins assigned to *Cebus imitator*, we mapped each protein group to orthologs using eggNOG-mapper v2 (Cantalapiedra et al., 2021) with the precomputed eggNOG v5.0 clusters and phylogenies (Huerta-Cepas et al., 2019).

Protein groups not found in at least 50% of samples were removed from downstream analysis. Three samples were removed from the dataset after failing quality control, resulting in a total of 45 samples from 24 individuals.

### Statistical analysis

#### Enrichment analyses

We identified the cellular origins of all *C. imitator* proteins by using a cell-type-specific enrichment analysis (CSEA) of all proteins found in at least 50% of samples. We used WebCSEA (Dai et al., 2022) to search the list of *C. imitator* gene (protein) IDs against 1355 human tissue-cell type panels to identify patterns of cellular enrichment. Significance was calculated by ranking permuted p-values across all categories which were then combined into overall p-values using Fisher’s method and Bonferroni correction (Dai et al., 2022). Gene set enrichment analyses were conducted with gProfiler2 (Kolberg et al., 2020). Cebus candidate proteins were searched for enrichment in the the following databases: Gene Ontology Biological Processes (GO:BP) and Molecular Functions (GO:MF), Reactome, the Comprehensive Resource of Mammalian Protein Complexes (CORUM), and the Kyoto Encyclopedia of Genes and Genomes (KEGG).

#### Ecological predictors of protein intensity

For each proteome of ecological interest, we calculated mean normalized intensity values within individuals across all protein groups found in at least 50% of samples. From these values we calculated the average intensity of fruit, invertebrate, and parasite proteins found in each sample. In the case of the dietary invertebrates, protein groups from multiple species were identified, so we averaged the intensities across all species to yield a single dietary invertebrate protein intensity value.

Similarly, we calculated average protein abundances for the multispecies genera *Bifidobacterium* and *Streptococcus*, which dominate the gut of *C. imitator* at SSR.

We built linear mixed effects models with the lmer function using the lme4 (Bates et al., 2015) and lmerTest (Kuznetsova et al., 2017) packages in R (v4.2.2) to predict the intensity of each *C. imitator* protein group. Fixed effects included age, sex, season, mean normalized intensity of dietary fruit, and mean normalized intensity of dietary invertebrates. Individual identity was included in each model as a random effect. For each model, age was measured as the log transformed number of days since an individual’s birth. However, because ageing effects can be nonlinear and especially apparent in aged individuals, we also reprocessed our models with age as a categorical value (young adult: <16 years; old adult: >16 years) based on the age distribution in our data set. The significance of individual predictors was tested in the models using age as a continuous variable. To avoid duplication, the models using age as a categorical value were only used to test for categorical age as an additional predictor. For each protein model, we ran an ANOVA and extracted fixed effect coefficients to calculate the log2 fold difference for each predictor, and set a cutoff of one for a meaningful difference in effect size.

P-value alpha levels for the significance of individual predictors were set at 0.05 and FDR corrections were secondarily calculated. To account for correlated predictors, we generated variance inflation factors (VIF) for each model using the vif command in the package car (Fox & Weisberg, 2019). Models with a VIF above 5 for any predictor were excluded from further analysis.

Subsequently, we built linear models using the same approach to predict the average intensity of non-host proteins found in the *C. imitator* gut. Models were constructed to predict the protein intensity of *Bifidobacterium*, *Streptococcus*, and *Strongyloides* using season, age, sex, and the protein intensities of fruit and invertebrates as fixed effects. Individual identity was included as a random effect. Models to predict the protein intensity of fruit and invertebrates used season, age, and sex as fixed effects and individual as a random effect. Additionally, we ran wilcoxon tests for significance simple comparisons between groups depicted in boxplots. All models are presented in table S2.

## Results

### Fecal proteome contents and enrichment

We identified 1143 protein groups represented in 67 of the 72 species in our proteome database. Of these, 1051 were assigned uniquely to single proteins, and 92 were razor peptides (Table S3). Among the broader categories of interest, we identified 394 protein groups from *Cebus imitator*, 491 from gut microbes, 153 from dietary fruit, 91 from dietary invertebrates, and 14 from intestinal parasites. As is common with DDA analysis of heterogeneous samples, we observed a substantial number of missing intensity values for individual protein groups. When only considering those found in at least 50% of samples, we identified 182 protein groups (161 unique; 21 razor) from 14 species: 154 protein groups from *C. imitator*; 15 from gut microbes; four from dietary fruit; eight from dietary invertebrates; and one from an intestinal parasite.

We classified the cellular composition of the *C. imitator* fecal proteome by searching the set of 154 reliable *C. imitator* proteins for enrichment in 1355 human-tissue cell types (TCs) using WebCSEA (Dai et al., 2022). 32 TCs were significantly enriched at the Bonferroni corrected p-value of 3.69x10^-5^ (Figure 1, Table S4). These TCs belonged to nine general cell types: enterocytes, goblet cells, epithelial cells, acinar cells, pancreas exocrine glandular cells, hepatocytes, neutrophils, endothelial cells, and kidney loop of Henle epithelial cells. The TCs corresponded to eight organ systems, but were dominated by cell types belonging to the digestive system (19 TCs).

**Figure 1:**
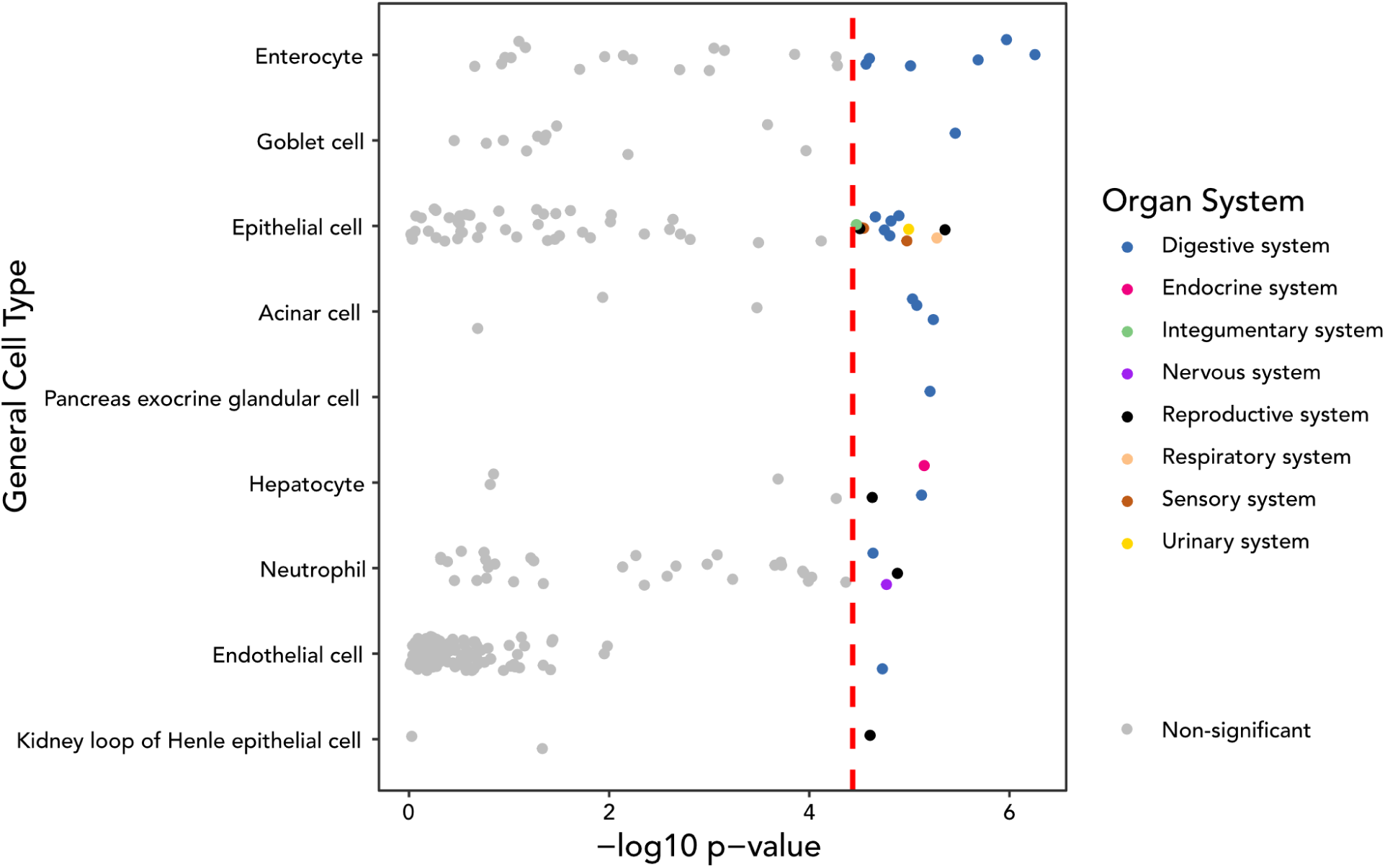
Cellular origins of fecally derived *Cebus imitator* proteins. Each point corresponds to one of 1355 human tissue-cell types from the WebCSEA human scRNA-seq tissue panel collection, which have been categorized into general cell types along the y-axis. X-axis values correspond to the -log10 combined p-value for tissue-cell-type specificity. The Bonferroni-corrected (p = 3.69x10^-5^) threshold is depicted with the red dashed line.

Gene set enrichment analysis (GSEA) with gProfiler2 (Kolberg et al., 2020) revealed 302 functionally enriched terms in the *C. imitator* fecal proteome: 195 GO Biological Processes, 69 GO Molecular Functions, 19 Kegg pathways, 28 Reactome pathways, and 9 CORUM protein complexes (Table S5). Across these databases, enriched terms were dominated by functions related to digestion and immune function (Table 1). Given the high number of enriched GO biological processes, we generated a similarity matrix of GO:BP terms using simplifyGO (Gu & Hübschmann, 2023). Matrix similarity values were then used to cluster GO terms into 12 clusters with summarized terminology (Figure 2). Clustering of GO:BP terms indicated a prioritization of similar processes, such as immune response, catabolic processes, blood circulation, and other metabolic processes related to digestion and homeostasis.

**Figure 2:**
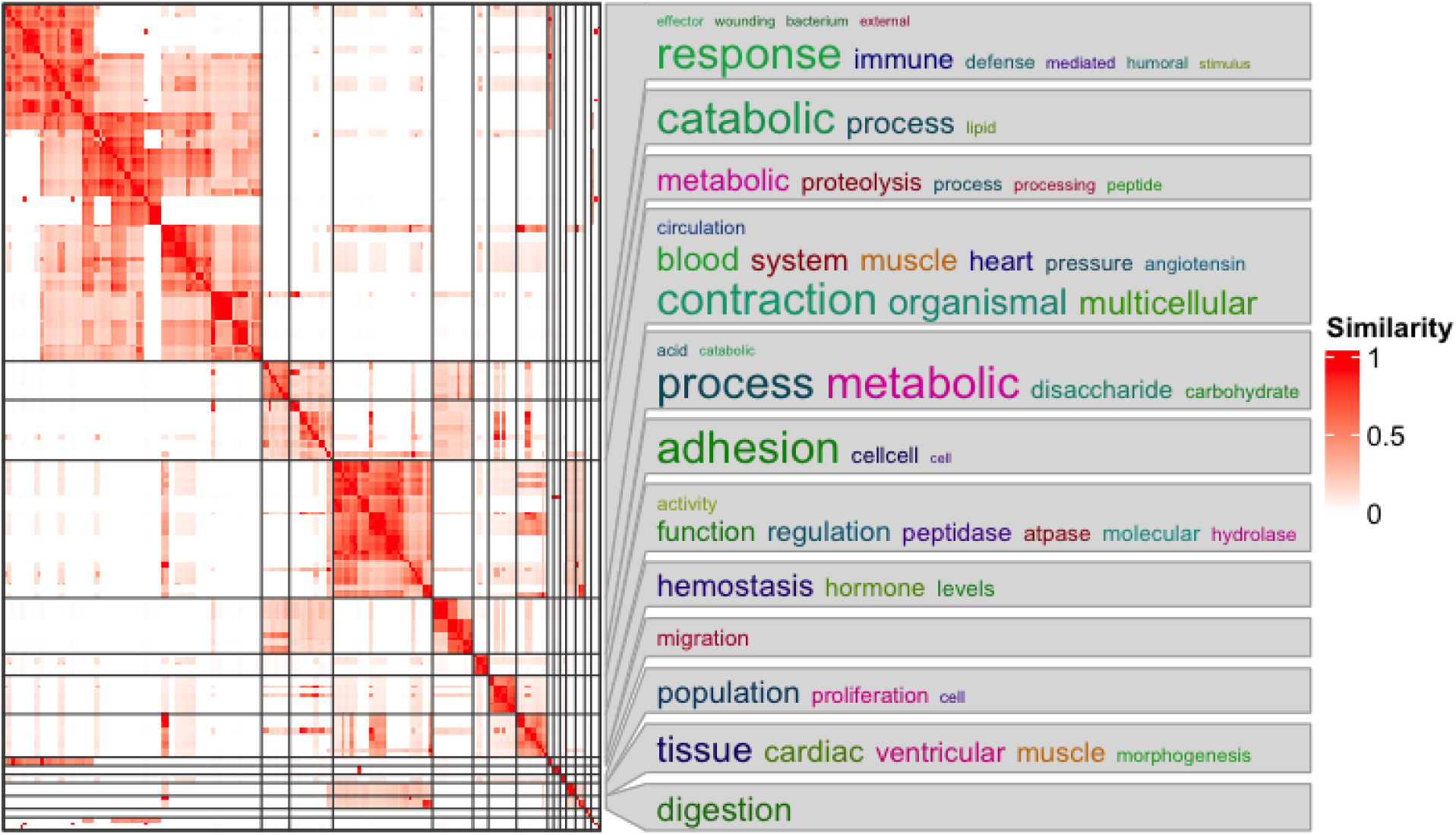
Clustering of 192 GO Biological Process terms with simplifyGO. The matrix heatmap on the left depicts the similarity scores between pairs of GO terms, which have been clustered into 12 groups. The word cloud on the right summarizes the terminology (scaled by frequency) in each cluster.

**Table 1:** Top five enriched terms in the *C. imitator* fecal proteome from KEGG, Reactome, Gene Ontology (Biological Processes and Molecular Functions), and CORUM.

| Database | Term ID | Term | P adjusted |
| --- | --- | --- | --- |
| KEGG | KEGG:04972 | Pancreatic secretion | 2.45E-11 |
| KEGG | KEGG:04975 | Fat digestion and absorption | 1.44E-06 |
| KEGG | KEGG:04974 | Protein digestion and absorption | 2.81E-06 |
| KEGG | KEGG:00010 | Glycolysis / Gluconeogenesis | 1.26E-05 |
| KEGG | KEGG:01100 | Metabolic pathways | 4.42E-05 |
| REAC | REAC:R-HSA-6798695 | Neutrophil degranulation | 1.19E-43 |
| REAC | REAC:R-HSA-168249 | Innate Immune System | 5.41E-32 |
| REAC | REAC:R-HSA-168256 | Immune System | 2.50E-23 |
| REAC | REAC:R-HSA-8935690 | Digestion | 9.33E-13 |
| REAC | REAC:R-HSA-8963743 | Digestion and absorption | 1.30E-11 |
| GO:BP | GO:0006952 | defense response | 5.90E-24 |
| GO:BP | GO:0002376 | immune system process | 9.26E-22 |
| GO:BP | GO:0044419 | biological process involved in interspecies interaction between organisms | 1.27E-20 |
| GO:BP | GO:0009056 | catabolic process | 6.30E-20 |
| GO:BP | GO:0019730 | antimicrobial humoral response | 1.04E-19 |
| GO:MF | GO:0016787 | hydrolase activity | 4.24E-21 |
| GO:MF | GO:0008233 | peptidase activity | 2.37E-19 |
| GO:MF | GO:0061134 | peptidase regulator activity | 6.32E-19 |
| GO:MF | GO:0008238 | exopeptidase activity | 6.66E-19 |
| GO:MF | GO:0004866 | endopeptidase inhibitor activity | 3.23E-18 |
| CORUM | CORUM:6823 | CP-LF-MPO complex | 6.19E-04 |
| CORUM | CORUM:6135 | Ferritin complex | 4.98E-02 |
| CORUM | CORUM:6821 | TTR-RBP complex | 4.98E-02 |
| CORUM | CORUM:426 | Meprin A | 4.98E-02 |
| CORUM | CORUM:6824 | CP-LF complex | 4.98E-02 |

### Quantification of non-host proteins

In total, we identified proteins assigned to 66 of 71 non-*Cebus* species in our protein database, including dietary fruits (1/1), dietary invertebrates (6/6), microbes (57/62), and gut parasites (2/2). However, for deeper analysis we focussed on those taxa consistently present in the gut of *C. imitator*. After filtering our protein intensity data to include only those protein groups found in at least 50% of samples (Figure S1), we identified 12 species of interest, including six bacteria known to inhabit the gut of *C. imitator* (*Bifidobacterium lemurum*, *Bifidobacterium scardovii*, *Blautia argi*, *Enterococcus thailandicus*, *Streptococcus ferus*, and *Streptococcus himalayensis*), one dietary fruit proxy (*Ficus carica*), four invertebrate prey proxies (*Heliothis virescens*, *Nezara viridula*, *Trachymyrmex cornetzi*, and *Vespula vulgaris*) and one intestinal parasite proxy (*Strongyloides venezuelensis*).

In order to generate a general intensity level for categories of ecological interest, we averaged the normalized protein intensities across each individual for dietary invertebrates, dietary fruit, *Bifidobacterium*, and *Streptococcus* (Table S6). Out of 45 samples, we were able to quantify protein intensity for dietary fruit in 42, dietary invertebrates in 41, *Bifidobacterium* in 40, *Streptococcus* in 39, and *Strongyloides* in 26. We observed a significantly higher level of normalized mean protein intensity for invertebrates than fruit (figures 3A, S2) in wilcoxon tests (W=556, p=0.005). Fruit intensity was also significantly elevated in the wet season samples (W = 135, p= 0.032; F=4.942, p=0.032) and older individuals (F=5.436, p=0.025) (Figure S3). Invertebrate protein intensity was not significantly associated with seasonality (W=141, p=0.078; F=3.399; p=0.082) or other linear model predictors. We also did not observe a significant association between the presence of fruit or invertebrates in feces (as recorded during collection) and protein intensity; however only three samples lacked visual evidence of fruit. We did not detect a significant association for *Bifidobacterium* or *Strongyloides* in our linear models, but *Streptococcus* protein intensity was significantly predicted by fruit intensity (F=6.367, p=0.024), invertebrate intensity (F=11.239, p=0.002), and season (F=12.1425, p=0.002).

**Figure 3:**
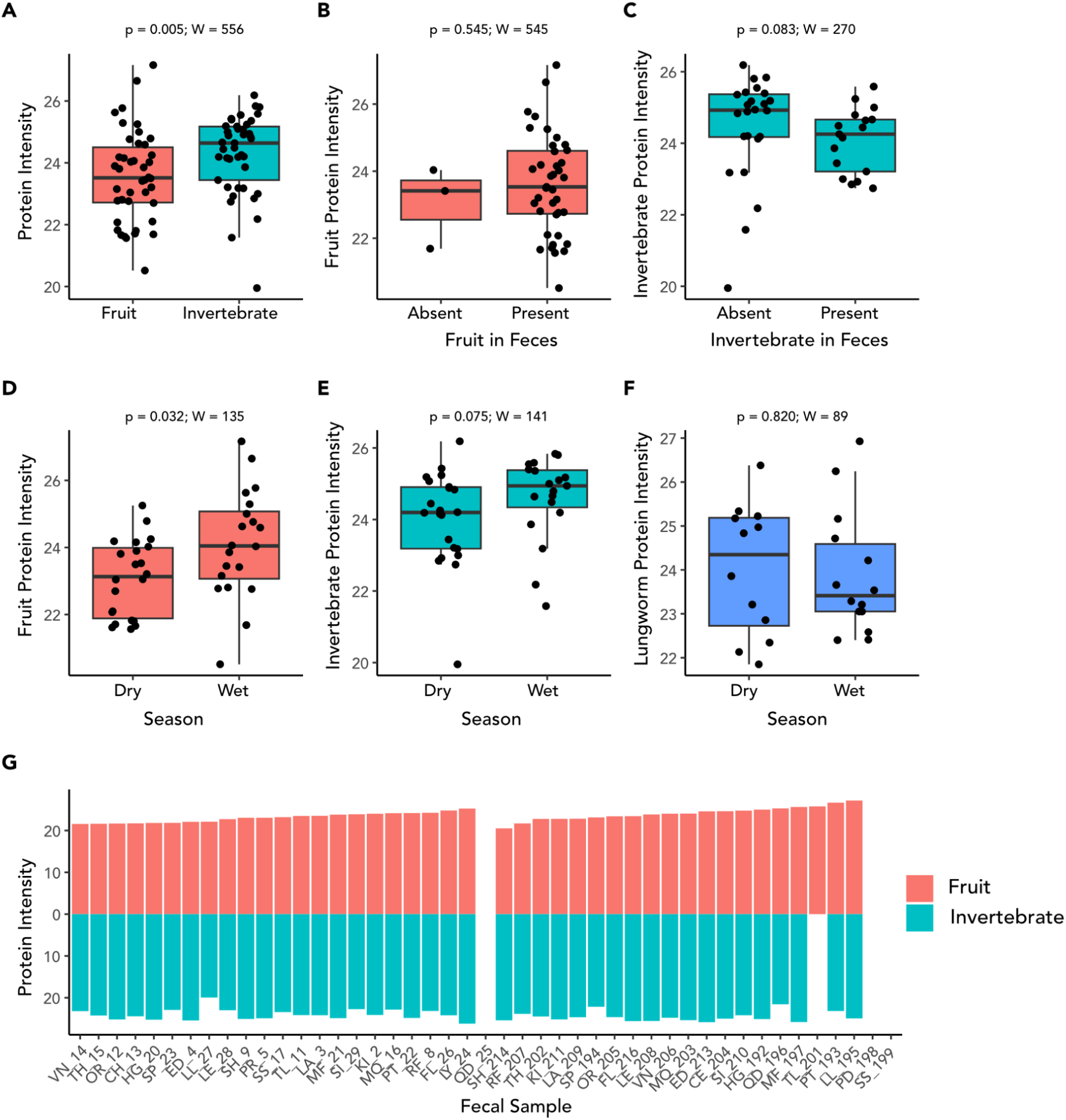
Normalized intensities of fruit and invertebrate proteins in feces of *C. imitator*. A) Across the entire dataset, invertebrate protein intensity is significantly higher than fruit intensity. B) There is no significant difference in fruit intensity for samples with/out visibly detectable fruit. C) There is no significant difference in invertebrate intensity for samples with/out visibly detectable Invertebrates. D) Fruit protein intensity is significantly higher in the wet season. E) Invertebrate protein intensity is not significantly higher in the wet season. F) There is no significant difference in lungworm intensity across seasons. G) Barplots showing the normalized protein intensity levels for fruit and invertebrates in each capuchin fecal sample. White bars for four samples (QD_25, TL_201, PD_198, and SS_199) indicate missing data.

### Differential abundance of *Cebus imitator* proteins

To identify which ecological predictors could be influencing the abundance of proteins in the *C. imitator* proteome, we built linear mixed effects models to predict the intensity of each protein assigned to *C. imitator* that was found in at least 50% of individuals. Fixed effects included age (continuous or categorical), sex, dietary fruit intensity, and dietary invertebrate intensity. We controlled for replicate sampling by setting the individual and sampling bout (wet or dry season) as random effects. Given the small sample size and complexity of the model structure, we set low thresholds for identifying candidate proteins: log2 fold change greater than +/- 1 and an uncorrected alpha level of p < 0.05. From the 154 protein groups observable in at least 50% of our samples, we identified 14 candidate proteins associated with age (APOD, CPO, DEFA4, EPX, LCT, S100A9, TFF2, and an immunoglobin V-type protein), sex (IGKV2D-29, IGLL1, SERPINC1), and the abundance of dietary insect proteins in the gut (CHIA, PLSCR1, and XPNPEP2). We also identified 27 proteins associated with seasonal differences, including ANXA2, CHIA, CLCA1, CLPS, CSTA, EPX, FCGBP, GSR, IGHE, IGHV3-49, IGLV8-61, LAP3, LCT, ORM1, PEPD, PLB1, PLSCR1, PNLIP, PNLIPRP1, PRSS2, SERPINA1, SERPINA3, SERPING1, XPNPEP2, ZG16, ZG16B, and one basic proline rich protein. (Figure 4, Table S7). Given the relatively large number of proteins associated with seasonal sampling period, we searched this gene list for enrichment in ontology and pathway databases. We identified 30 enriched categories (Table S8), including acute inflammatory response (GO:0006953), innate immune system (REAC:R-HSA-168249), neutrophil degranulation (REAC:R-HSA-6798695), fat digestion and absorption (KEGG:04975), vitamin digestion and absorption (KEGG:04977), and digestion of dietary carbohydrate (REAC:R-HSA-189085).

**Figure 4:**
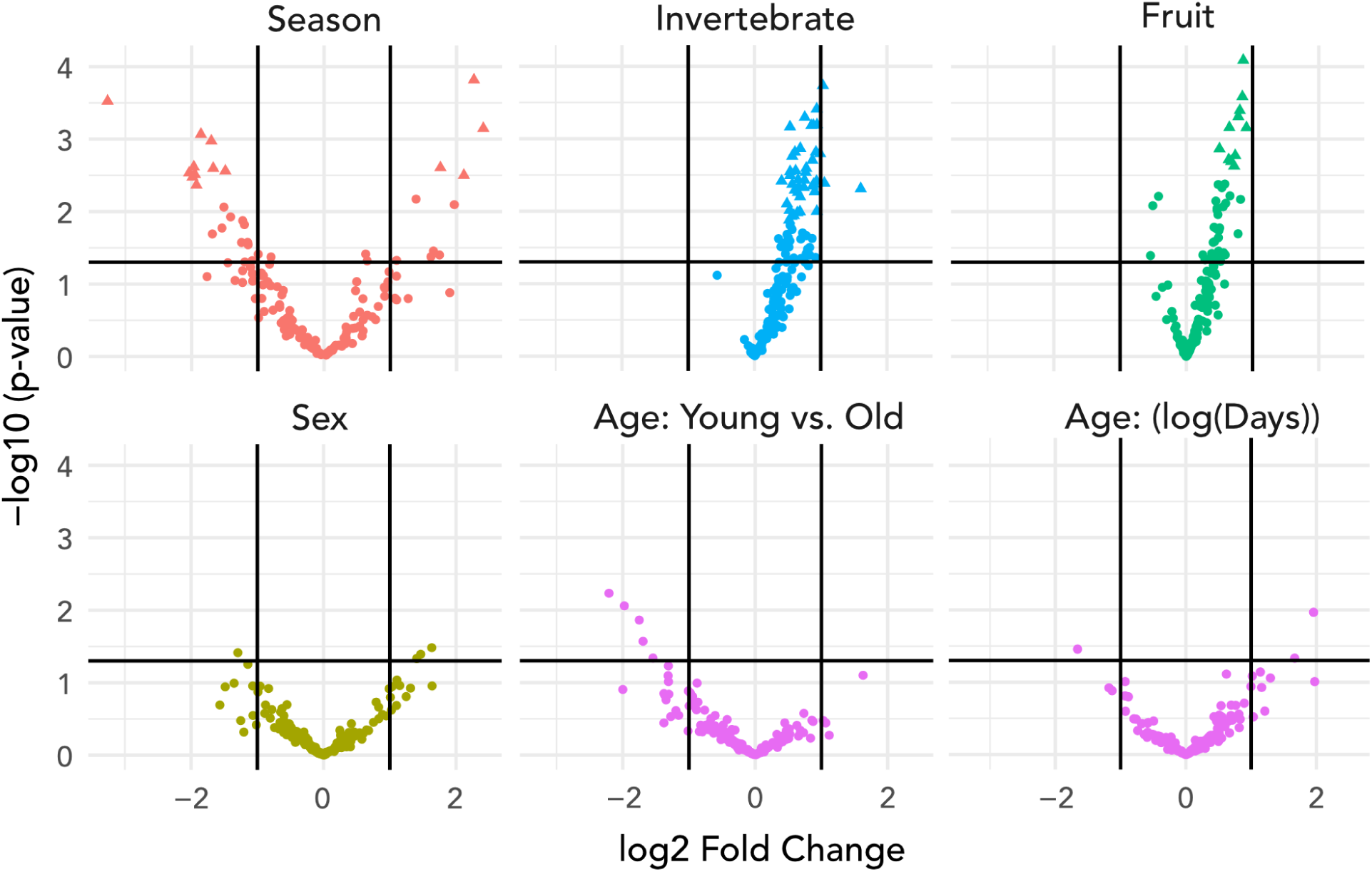
Volcano plots of predictors of differential abundance of *C. imitator* proteins from linear mixed effects models, including sampling season, the normalized protein intensity of dietary invertebrates, the normalized protein intensity of dietary fruit, individual sex, and individual age (both as continuous and categorical variables). The horizontal line marks the uncorrected p-value of 0.05. Triangles indicate proteins that pass the FDR cutoff for a given predictor, and circles do not. The vertical lines correspond to a log2 fold difference of +/- 1.

## Discussion

We are the first to apply quantitative fecal proteomics to study the ecology of a free-ranging population of mammals. We generated direct measures of differential protein abundances from both host and exogenous sources in the white faced capuchin gut that can be integrated with long-term field research on the SSR population to explore the complex interaction between the seasonal dry forest biome, dietary ecology, and host physiology. With this new approach, we have been able non-invasively to: 1) categorize the cellular composition and functional enrichment of the capuchin gut; 2) reveal how host cellular physiology is influenced by host-specific traits and ecological variation; and 3) quantify the abundances of consumed dietary items and multicellular parasites. We suggest that fecal proteomics is a viable method for the molecular ecology of free-ranging mammals and can offer substantial benefits over traditional metagenomics.

We identified 394 proteins from the *C. imitator* gut and localized the most abundant proteins to cell types and organ systems, which demonstrates the tissue specificity of the information generated by fecal proteomics. This is a major advance over traditional metagenomics, for which it remains difficult to generate direct quantitative information about host cellular physiology non-invasively. Metagenomic approaches to understanding gut homeostasis largely rely on microbial information–taxonomy, gene families, and pathways–that can be used as proxy information for host gut health. Further, DNA sequencing data can only identify the functional capacity of the gut and cannot provide information on the gene or protein content of the host organism that is specific to the tissues of interest. As we have demonstrated, fecal proteomics allows for the measurement of cell-specific proteins from the host organism.

These measures provide direct, real-time evidence of host physiology and health status in individual capuchins.

We identified a total of 41 candidate proteins with differentially abundant intensities associated with season, age, diet, or sex. Among these proteins, the 27 associated with seasonal differences represented by far the largest proportion, and enrichment analysis indicates that these proteins are overrepresented in pathways associated with digestion, inflammation, and immune activity. This clustering is consistent with the pattern of seasonal challenges at the SSR tropical dry forest biome, where resident primates must survive extended periods of drought, dietary fruit scarcity, loss of muscle mass, and increased gut parasite diversity (Bergstrom et al., 2017; Campos et al., 2020; Pinto et al., 2023). Furthermore, previous research on the SSR capuchins has demonstrated selection on genes involved in the metabolism of carbohydrates and lipids and muscular wasting (Orkin, Montague, et al., 2021).

Given these observations, the seasonal differential abundance of ORM1 is particularly interesting. ORM1 is known to regulate injury-induced angiogenesis and inflammation (Ligresti et al., 2012), but it is also involved in the maintenance of muscular strength and endurance in age-related sarcopenia (Altab et al., 2024), suggesting that it could play an important role in how the SSR capuchins compensate for seasonal challenges across their lifespans. However, we caution that the proteins identified as differentially abundant across seasons should be treated as candidates for further analysis. Although samples were drawn from 24 individuals, the seasonality comparison is made across only two timepoints. As such, other factors could have influenced the protein abundances we observed. Nonetheless, the discovery of candidate proteins involved in physiological processes that have been identified as under selection from genome scans demonstrates the promise of fecal proteomics to provide real-time measures of physiological status that can be integrated into an ecological and evolutionary framework.

Among the eight proteins associated with age in our differential abundance list, two of them (APOD and S100A9) are listed in the GenAge database as being significantly overexpressed in healthy ageing microarray datasets (Tacutu et al., 2018). We observed a significant increase in APOD intensity with age as a continuous variable (Figure 5A). APOD has been heavily studied in the context of human ageing research, particularly for its association with Alzheimer’s and other neurodegenerative disorders (Muffat & Walker, 2010). It is widely expressed in the body, playing key roles in lipid metabolism and oxidative stress, extending lifespan (de Magalhães et al., 2009; Muffat et al., 2008; Perdomo & Henry Dong, 2009). In a metaanalysis of genes consistently overexpressed in ageing individuals, APOD had the strongest association of all 56 genes examined (de Magalhães et al., 2009). S100A9 proteins are primarily derived from immune cells and upregulated during stress and other inflammatory processes (Wang et al., 2018). It is strongly associated with age-related inflammation and shifting abundances of the S100A8-S100A9 heterocomplex (calgranulin) have been reported as a feature of intrinsic ageing in a wide range of mammalian tissue (Swindell et al., 2013).

**Figure 5:**
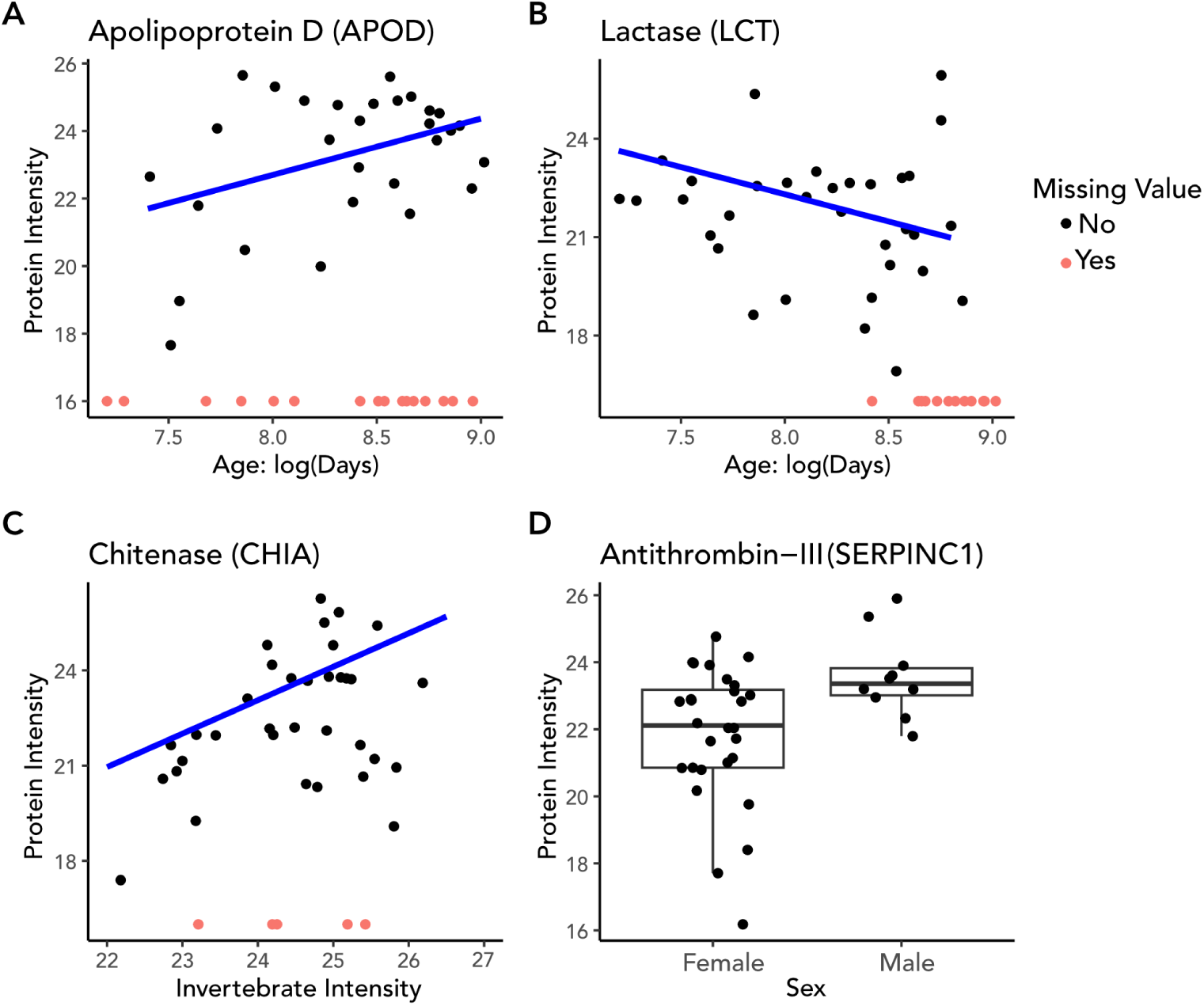
Intensities of *C. imitator* proteins identified in the fecal proteome and their associated predictors. All samples with missing intensity values are depicted in red for visualization. Trendlines are plotted from the marginal predictions of the mixed models. A) Apolipoprotein D increases with age. B) Lactase is persistent through adulthood in *C. imitator* but decreases with age. C) Chitinase increases with the amount of dietary invertebrates in the gut of *C. imitator*. D) Antithrombin-III is significantly lower in females than males.

We also observed several proteins that are differentially abundant in males and females, but for the most part their functional roles are less clearly related to differences between the sexes. However, sex differences in thrombosis have been observed for SERPINC1 in laboratory experiments on mice (Wong et al., 2008). SERPINC1 encodes antithrombin III, which regulates the blood coagulation cascade, and experiments demonstrate that SERPINC1 and other coagulation factors expressed in the liver are heavily influenced by growth hormone secretion, which is strongly biased by sex (Wong et al., 2008). We observed a significantly higher level of SERPINC1 intensity in males from SSR (Figure 5D), although we caution that this could be related to differences in sample size. Capuchins at SSR, especially males, have been frequently observed recovering from major wounds, and future research could target questions about sex biases in the acquisition and healing of wounds as it relates to variation in SERPINC1.

Curiously, SERPINC1 has also been observed to be under selection at the genomic level in Costa Rican *C. imitator*, when comparing wet and dry forest populations (including SSR) (Orkin, Montague, et al., 2021). While it is not clear how this pattern of selection would be related to sex differences, it does suggest that SERPINC1 plays an important physiological role in the SSR capuchins that should be investigated further.

Until now, it has not been possible to quantify the amount of protein in a fecal sample that is derived from particular dietary items. Previous research has relied on visible evidence of particular foods, such as insect wings or fig seeds, in feces to classify the presence or absence of fruit or invertebrate prey. However, given the uncertain digestive efficiency of capuchins (or other species) it is not *a priori* evident if visual indicators should be treated as quantifiable measures of dietary composition. Likewise, despite there being a substantial difference in monthly fruit biomass at SSR during our two sampling seasons, nearly all fecal samples had both visible evidence of fruit and substantial fruit protein intensities (Figure 3). As such, it is not surprising that we did not find a significant association between the presence of fruit or insect remains and their protein intensities. In fact, our ability to successfully correlate the abundance of a digestive enzyme (CHIA) with the abundance of invertebrate proteins in feces strongly supports the accuracy of our quantitative approach to dietary ecology. Chitinase plays a critical role in the digestion of arthropod exoskeletons and the cell walls of fungi (Janiak, 2016). As such, it is not inherently surprising to find an association between invertebrates composed of chitin and the enzyme that degrades it. However, it does serve as an important validation of our ability to quantify the abundance of broad categories of dietary items using homologous proteins from non-target species. While the sampling season also predicted CHIA intensity, this is likely a result of the higher amount of insects found in the wet season, which was nearly significant (Figure 3E).

Strikingly, we observed a high frequency and quantity of lactase (LCT) proteins assigned to *C. imitator* in the recovered proteins from individuals of all ages. This is highly unexpected, given that humans are the only primates known to have evolved lactase persistence (Janiak, 2016). The demonstration of lactase persistence in *C. imitator* would be a startling result, and as such it needs direct validation before the claim can be made. Nonetheless, there is some evidence to suggest its plausibility. First, *C. imitator* is notable for the very high abundance of *Bifidobacterium* in its gut microbiome, despite the lack of evidence for post-weaning milk consumption (Orkin, Campos, et al., 2019), after which *Bifidobacterium* abundances typically decline (Tsutaya et al., 2021). One possible explanation is that while both LCT from *C. imitator* and the similar beta-galactosidase (lacZ) enzyme in *Bifidobacterium* are capable of digesting lactase (He et al., 2005), in the adult *C. imitator* gut they are involved in the digestion of non-lactose sugars. While it is possible that spectra belonging to lacZ were incorrectly assigned to LCT, this would be surprising. Our metaproteomic database included 128 beta-galactosidase sequences from a wide range of organisms, including those from *B. scardovii, B. lemurum*, and *B. catenulatum*, and our 479 spectral matches were assigned to 28 peptides of LCT from *C. imitator*, with a top peptide probability of 0.999, making such an error highly unlikely. Furthermore, Tsutaya et al. (2021) demonstrated the loss of spectral matches for lactase protein groups in macaques post weaning. Thus, it stands to reason that proteomic approaches should be able to reliably distinguish primate LCT from exogenous beta-galactosidase. It is also worth noting that unlike other proteins in our data set, all the undetected values for LCT come from the oldest individuals, as opposed to being spread randomly across capuchins of all ages (Figure 5B), which could suggest an age-dependent effect of lactase persistence. Nonetheless we urge caution in the interpretation of this result, which will require further analysis and validation.

We detected a strong signal of *Bifidobacterium* and *Streptococcus* in the *C. imitator* gut microbiome, which fits expectations from previous microbiome research (Orkin, Campos, et al., 2019; Orkin, Webb, et al., 2019). *Streptococcus* intensity was significantly predicted by fruit intensity, invertebrate intensity, and season in our protein data. Elsewhere, metagenomic data has also indicated that *Streptococcus* abundance is correlated with the presence of insects in *C. imitator* feces at SSR, which is consistent with the results obtained here (Orkin, Campos, et al., 2019). However, while *Bifidobacterium* abundance has previously been associated with fruit abundance in this population (Orkin, Campos, et al., 2019), we were unable to demonstrate the same finding, likely as a result of low power, given the presence of fruit in nearly all samples used in the present study (Figure 3B). High parasite loads have also been observed in SSR capuchins, including the lungworm genus *Stronglyoides* and *Giardia* (Parr et al., 2013; Pinto et al., 2023). We were able to quantify proteins from *Strongyloides* and *Giardia* in 26 and 4 samples, respectively, but did not identify any significant predictor variables. Nonetheless, these results provide promising data on parasite presence and abundance. Future work could search for the presence of tissue specific proteins in parasites to distinguish adults from larvae and eggs for quantification.

Despite the advances we have demonstrated, challenges for methodological development in fecal proteomics remain. A limitation of our approach was the high number of missing intensity values spread across our dataset, which limited the power of our statistical analysis. This pattern is likely the result of using a data-dependent acquisition (DDA) mass spectrometry protocol. DDA selects the most intense precursor ions during the MS1 phase for fragmentation in the MS2 stage. While this provides a targeted analysis of the most abundant proteins across samples, a large number of low intensity ions are not analyzed. For instance, the fact that we only observed haptoglobin in four samples does not indicate its absence in the majority of samples if it is present in low abundance. In the case of highly complex mixtures like feces, newly developed data-independent acquisition (DIA) protocols that sample all fragment ions within defined mass-to-charge windows are likely to provide more consistent and representative quantification of the fecal proteome. Given that only a single well-defined peptide is necessary for protein quantification, future studies could also use a parallel-reaction monitoring (PRM) approach for targeted proteomics. With further development of spectral libraries for *C. imitator* (or any species of interest) it will become possible to select precise spectral windows that coincide with peptides/proteins of interest, allowing for highly accurate label-free quantification from feces.

We have quantified the intensity of proteins from host-specific and exogenous sources in the gut of white-faced capuchins from a free-ranging population using non-invasively collected fecal samples. Proteins assigned to *C. imitator* were strongly localized to cells in the gut lining and predominantly involved in digestive and immune functions. We further demonstrated how the quantity of individual *C. imitator* proteins differs in response to variation in season, diet, age and sex. Our ability to identify differentially abundant intensities of proteins in the host gut that are associated with these innate and external variables is a major contribution to ecological proteomics. Additionally, we were able to provide quantitative measures of dietary items, multicellular parasites, and gut microbes, providing an advance over standard metagenomic techniques. We have also overcome the challenge of limited proteomic resources for database construction by relying on protein homology from closely related species to assign and quantify proteins into categories of ecological interest. Together, these results demonstrate the utility of fecal proteomics for the study of free-ranging primates and other mammals.

## Data accessibility

At the time of publication, mass spectral RAW files will be deposited in ProteomeXchange and code will be made available in a repository on github.

## Acknowledgements

We are grateful to Gwen Duytschaever for assistance with laboratory work and Luiz Gustavo Nohueria de Almeida for statistical advice. We also thank Roger Blanco, Maria Marta, Linda M Fedigan, and all members of the Santa Rosa field team.

## Supplemental Tables

**Table S1:** List of 72 species used for Fragpipe database search

**Table S2:** Full output of all protein models

**Table S3**: Details of identified protein groups

**Table S4**: Cell Set Enrichment Analysis (CSEA) results

**Table S5**: Gene Set enrichment of significant CSEA proteins

**Table S6**: Normalized intensity data from non-*Cebus* proteins used in models

**Table S7**: Differentially abundant proteins in *Cebus*

**Table S8**: GSEA results for seasonally enriched proteins.

---

**Figure S1:**
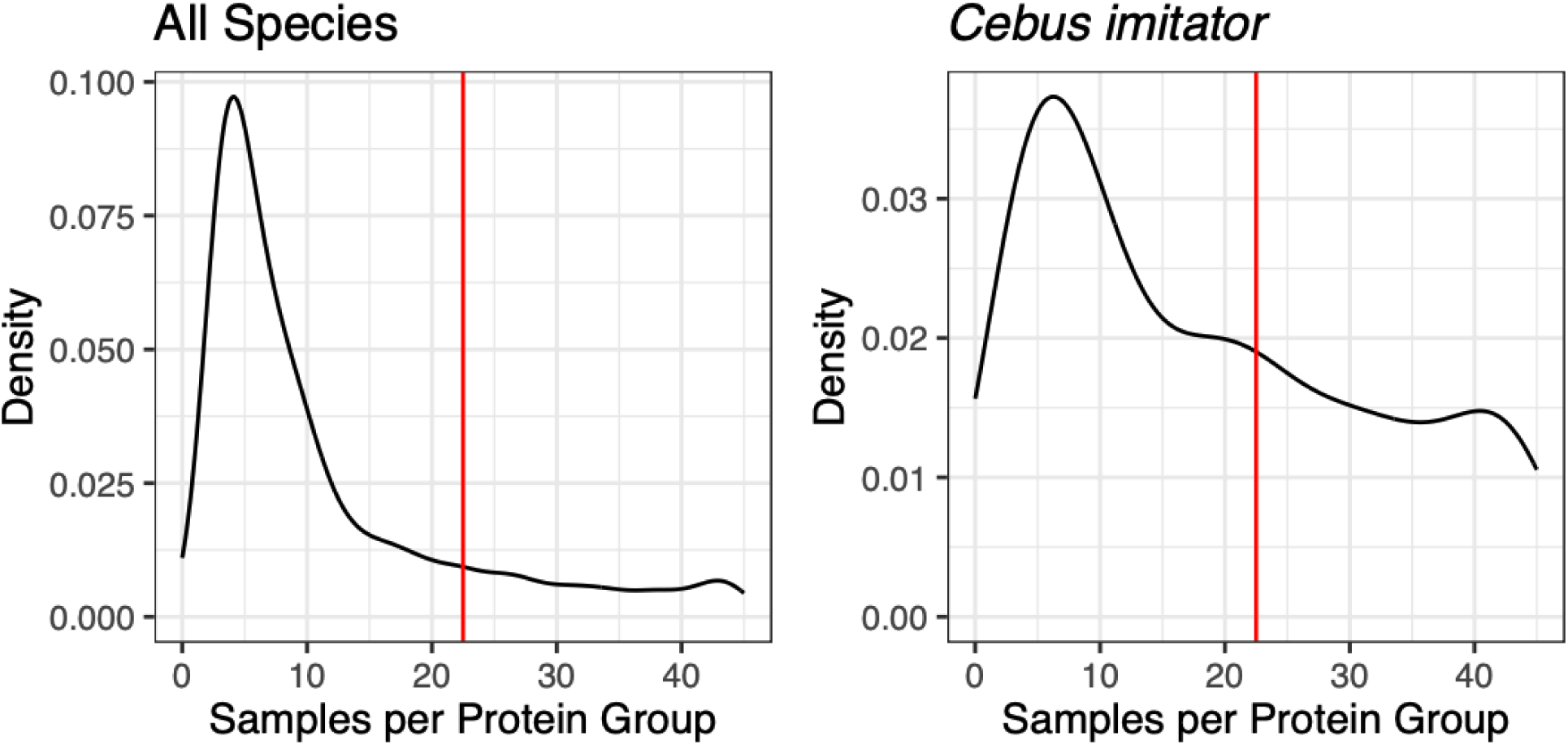
Density of protein groups per sample for all species (left) and *C. imitato*r only (right). Red line demarcates the presence of a protein group in 50% of samples.

**Figure S2:**
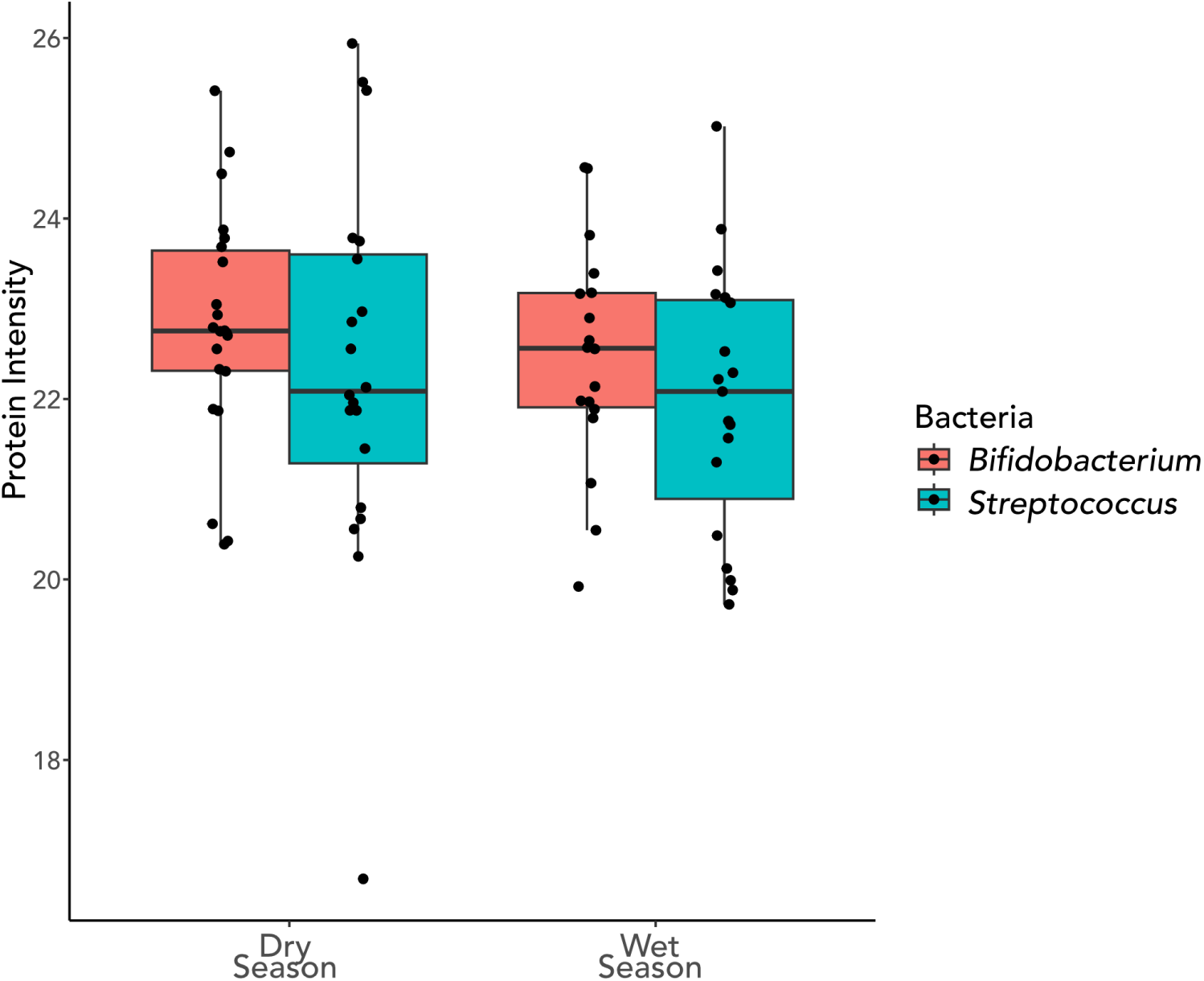
Mean normalized protein intensity of *Bifidobacterium* and *Streptococcus* in the dry and wet seasons at SSR.

**Figure S3:**
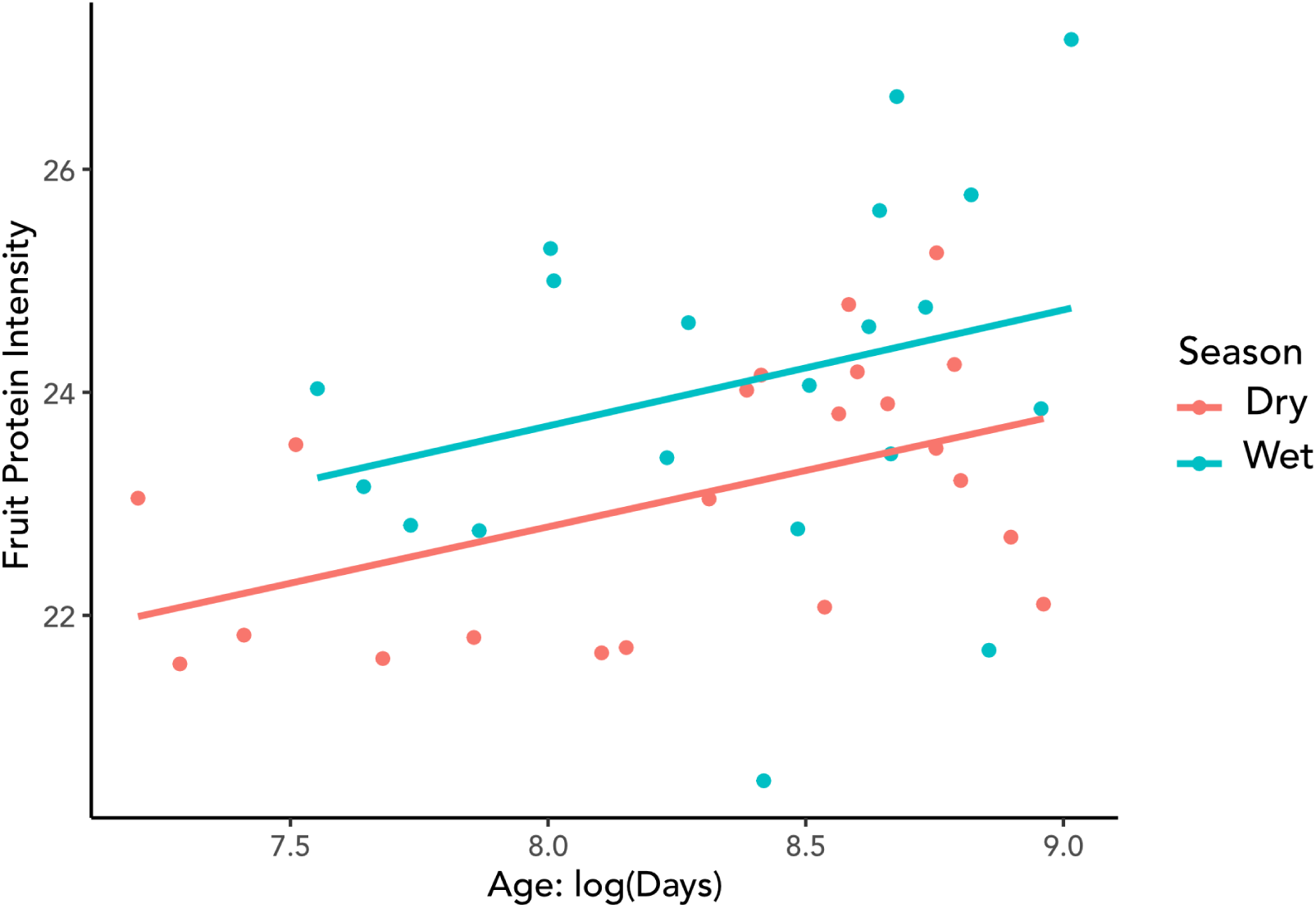
Mean normalized protein intensity of dietary fruit predicted by age. Regression lines and colors correspond to the dry and wet seasons.

## References

1. Altab, G., Merry, B. J., Beckett, C. W., Raina, P., Lopes, I., Goljanek-Whysall, K., & de Magalhães, J. P. (2024). Unravelling the transcriptomic symphony of sarcopenia: Key pathways and hub genes altered by muscle ageing and caloric restriction revealed by RNA sequencing. In bioRxiv (p. 2024.03.09.584213). 10.1101/2024.03.09.584213

2. Amato, K. R., G Sanders, J., Song, S. J., Nute, M., Metcalf, J. L., Thompson, L. R., Morton, J. T., Amir, A., J McKenzie, V., Humphrey, G., Gogul, G., Gaffney, J., L Baden, A., A O Britton, G., P Cuozzo, F., Di Fiore, A., J Dominy, N., L Goldberg, T., Gomez, A., … R Leigh, S. (2018). Evolutionary trends in host physiology outweigh dietary niche in structuring primate gut microbiomes. The ISME Journal. 10.1038/s41396-018-0175-0

3. Arandjelovic, M., & Vigilant, L. (2018). Non-invasive genetic censusing and monitoring of primate populations. American Journal of Primatology. 10.1002/ajp.22743

4. Bădescu, I., Katzenberg, M. A., Watts, D. P., & Sellen, D. W. (2017). A novel fecal stable isotope approach to determine the timing of age-related feeding transitions in wild infant chimpanzees. American Journal of Physical Anthropology, 162(2), 285–299.

5. Bates, D., Mächler, M., Bolker, B., & Walker, S. (2015). Fitting Linear Mixed-Effects Models Using lme4. *Journal of Statistical Software*, Articles, 67(1), 1–48.

6. Bergstrom, M. L., Emery Thompson, M., Melin, A. D., & Fedigan, L. M. (2017). Using urinary parameters to estimate seasonal variation in the physical condition of female white-faced capuchin monkeys (Cebus capucinus imitator). American Journal of Physical Anthropology, 163(4), 707–715.

7. Campos, F. A. (2018). A synthesis of long-term environmental change in Santa Rosa. In U. Kalbitzer & K. M. Jack (Eds.), Primate Life Histories, Sex Roles, and Adaptability - Essays in Honour of Linda M. Fedigan. Springer, New York.

8. Campos, F. A., & Fedigan, L. M. (2009). Behavioral adaptations to heat stress and water scarcity in white-faced capuchins (Cebus capucinus) in Santa Rosa National Park, Costa Rica. American Journal of Physical Anthropology, 138(1), 101–111.

9. Campos, F. A., Kalbitzer, U., Melin, A. D., Hogan, J. D., Cheves, S. E., Murillo-Chacon, E., Guadamuz, A., Myers, M. S., Schaffner, C. M., Jack, K. M., Aureli, F., & Fedigan, L. M. (2020). Differential impact of severe drought on infant mortality in two sympatric neotropical primates. Royal Society Open Science, 7(4), 200302.

10. Campos, F. A., Wikberg, E. C., Orkin, J. D., Park, Y., Snyder-Mackler, N., Cheves Hernandez, S., Lopez Navarro, R., Fedigan, L. M., Gurven, M., Higham, J. P., Jack, K. M., & Melin, A. D. (2024). Wild capuchin monkeys as a model system for investigating the social and ecological determinants of ageing. Philosophical Transactions of the Royal Society of London. Series B, Biological Sciences, 379(1916), 20230482.

11. Cantalapiedra, C. P., Hernández-Plaza, A., Letunic, I., Bork, P., & Huerta-Cepas, J. (2021). EggNOG-mapper v2: Functional annotation, orthology assignments, and domain prediction at the metagenomic scale. Molecular Biology and Evolution, 38(12), 5825–5829.

12. Chang, H.-Y., Kong, A. T., da Veiga Leprevost, F., Avtonomov, D. M., Haynes, S. E., & Nesvizhskii, A. I. (2020). Crystal-C: A computational tool for refinement of open search results. Journal of Proteome Research, 19(6), 2511–2515.

13. Dai, Y., Hu, R., Liu, A., Cho, K. S., Manuel, A. M., Li, X., Dong, X., Jia, P., & Zhao, Z. (2022). WebCSEA: web-based cell-type-specific enrichment analysis of genes. Nucleic Acids Research, 50(W1), W782–W790.

14. Dasari, M. R., Roche, K. E., Jansen, D., Anderson, J., Alberts, S. C., Tung, J., Gilbert, J. A., Blekhman, R., Mukherjee, S., & Archie, E. A. (2025). Social and environmental predictors of gut microbiome age in wild baboons. 10.7554/elife.102166.2

15. de Magalhães, J. P., Curado, J., & Church, G. M. (2009). Meta-analysis of age-related gene expression profiles identifies common signatures of aging. *Bioinformatics (Oxford*, England), 25(7), 875–881.

16. Fedigan, L., & Rose-Wiles, L. (1996). See how they grow: Tracking capuchin monkey populations in a regenerating Costa Rican dry forest. In M. A. Norconk, A. L. Rosenberger, & P. A. Garber (Eds.), Adaptive radiations of Neotropical primates (pp. 289–307). Springer.

17. Fox, J., & Weisberg, S. (2019). Functions to Accompany, An R Companion to Applied Regression (Third). Sage.

18. Fragaszy, D. M., Visalberghi, E., & Fedigan, L. M. (2004). The Complete Capuchin: The Biology of the Genus Cebus. Cambridge University Press.

19. Geiszler, D. J., Kong, A. T., Avtonomov, D. M., Yu, F., Leprevost, F. da V., & Nesvizhskii, A. I. (2021). PTM-Shepherd: Analysis and summarization of post-translational and chemical modifications from open search results. Molecular & Cellular Proteomics: MCP, 20, 100018.

20. Grieneisen, L., Dasari, M., Gould, T. J., Björk, J. R., Grenier, J.-C., Yotova, V., Jansen, D., Gottel, N., Gordon, J. B., Learn, N. H., Gesquiere, L. R., Wango, T. L., Mututua, R. S., Warutere, J. K., Siodi, L. ’ida, Gilbert, J. A., Barreiro, L. B., Alberts, S. C., Tung, J., … Blekhman, R. (2021). Gut microbiome heritability is nearly universal but environmentally contingent. Science, 373(6551), 181–186.

21. Gu, Z., & Hübschmann, D. (2023). SimplifyEnrichment: A bioconductor package for clustering and visualizing functional enrichment results. Genomics, Proteomics & Bioinformatics, 21(1), 190–202.

22. He, T., Priebe, M. G., Vonk, R. J., & Welling, G. W. (2005). Identification of bacteria with β-galactosidase activity in faeces from lactase non-persistent subjects. FEMS Microbiology Ecology, 54(3), 463–469.

23. Huerta-Cepas, J., Szklarczyk, D., Heller, D., Hernández-Plaza, A., Forslund, S. K., Cook, H., Mende, D. R., Letunic, I., Rattei, T., Jensen, L. J., von Mering, C., & Bork, P. (2019). eggNOG 5.0: a hierarchical, functionally and phylogenetically annotated orthology resource based on 5090 organisms and 2502 viruses. Nucleic Acids Research, 47(D1), D309–D314.

24. Janiak, M. C. (2016). Digestive enzymes of human and nonhuman primates. Evolutionary Anthropology, 25(5), 253–266.

25. Janzen, D. (2002). Tropical dry forest: area de Conservación Guanacaste, northwestern Costa Rica. In D. A. Perrow M (Ed.), Handbook of Ecological Restoration: Restoration in Practice (pp. 559–583). Cambridge University Press, Cambridge.

26. Kohler, D., Staniak, M., Tsai, T.-H., Huang, T., Shulman, N., Bernhardt, O. M., MacLean, B. X., Nesvizhskii, A. I., Reiter, L., Sabido, E., Choi, M., & Vitek, O. (2023). MSstats version 4.0: Statistical analyses of quantitative mass spectrometry-based proteomic experiments with chromatography-based quantification at scale. Journal of Proteome Research, 22(5), 1466–1482.

27. Kolberg, L., Raudvere, U., Kuzmin, I., Vilo, J., & Peterson, H. (2020). gprofiler2 -- an R package for gene list functional enrichment analysis and namespace conversion toolset g:Profiler. F1000Research, 9, 709.

28. Kong, A. T., Leprevost, F. V., Avtonomov, D. M., Mellacheruvu, D., & Nesvizhskii, A. I. (2017). MSFragger: ultrafast and comprehensive peptide identification in mass spectrometry-based proteomics. Nature Methods, 14(5), 513–520.

29. Kuznetsova, A., Brockhoff, P. B., & Christensen, R. H. B. (2017). LmerTest package: Tests in linear mixed effects models. Journal of Statistical Software, 82(13). 10.18637/jss.v082.i13

30. Lambert, J. E. (1998). Primate digestion: Interactions among anatomy, physiology, and feeding ecology. Evolutionary Anthropology, 7(1), 8–20.

31. Lambert, J. E., & Rothman, J. M. (2015). Fallback foods, optimal diets, and nutritional targets: Primate responses to varying food availability and quality. Annual Review of Anthropology, 44(1), 493–512.

32. Ligresti, G., Aplin, A. C., Dunn, B. E., Morishita, A., & Nicosia, R. F. (2012). The acute phase reactant orosomucoid-1 is a bimodal regulator of angiogenesis with time- and context-dependent inhibitory and stimulatory properties. PloS One, 7(8), e41387.

33. Ma, K., Vitek, O., & Nesvizhskii, A. I. (2012). A statistical model-building perspective to identification of MS/MS spectra with PeptideProphet. BMC Bioinformatics, 13 Suppl 16(S16), S1.

34. Matthews, J. K., Ridley, A., Kaplin, B. A., & Grueter, C. C. (2020). A comparison of fecal sampling and direct feeding observations for quantifying the diet of a frugivorous primate. Current Zoology, 66(4), 333–343.

35. Melin, A. D., Hogan, J. D., Campos, F. A., Wikberg, E., King-Bailey, G., Webb, S. E., Kalbitzer, U., Asensio, N., Murillo-Chacon, E., Cheves, H. S., Guadamuz, C. A., Schaffner, C., Kawamura, S., Aureli, F., Fedigan, L. M., & Jack, K. M. (2020). Primate life history, social dynamics, ecology, and conservation: contributions from long-term research in the Área de Conservación Guanacaste, Costa Rica. Biotropica. 10.1111/btp.12867

36. Moreno-Black, G. (1978). The use of scat samples in primate diet analysis. Primates; Journal of Primatology, 19(1), 215–221.

37. Muffat, J., & Walker, D. W. (2010). Apolipoprotein D: an overview of its role in aging and age-related diseases. *Cell Cycle (Georgetown*, Tex.), 9(2), 269–273.

38. Muffat, J., Walker, D. W., & Benzer, S. (2008). Human ApoD, an apolipoprotein up-regulated in neurodegenerative diseases, extends lifespan and increases stress resistance in Drosophila. Proceedings of the National Academy of Sciences of the United States of America, 105(19), 7088–7093.

39. Orkin, J. D., Campos, F. A., Myers, M. S., Cheves Hernandez, S. E., Guadamuz, A., & Melin, A. D. (2019). Seasonality of the gut microbiota of free-ranging white-faced capuchins in a tropical dry forest. The ISME Journal, 13(1), 183–196.

40. Orkin, J. D., Kuderna, L. F. K., & Marques-Bonet, T. (2021). The Diversity of Primates: From Biomedicine to Conservation Genomics. Annual Review of Animal Biosciences, 9, 103–124.

41. Orkin, J. D., Montague, M. J., Tejada-Martinez, D., de Manuel, M., Del Campo, J., Cheves Hernandez, S., Di Fiore, A., Fontsere, C., Hodgson, J. A., Janiak, M. C., Kuderna, L. F. K., Lizano, E., Martin, M. P., Niimura, Y., Perry, G. H., Valverde, C. S., Tang, J., Warren, W. C., de Magalhães, J. P., … Melin, A. D. (2021). The genomics of ecological flexibility, large brains, and long lives in capuchin monkeys revealed with fecalFACS. Proceedings of the National Academy of Sciences of the United States of America, 118(7), e2010632118.

42. Orkin, J. D., Webb, S. E., & Melin, A. D. (2019). Small to modest impact of social group on the gut microbiome of wild Costa Rican capuchins in a seasonal forest. *American Journal of Primatology*, e22985.

43. Parr, N. A., Fedigan, L. M., & Kutz, S. J. (2013). A coprological survey of parasites in white-faced capuchins (Cebus capucinus) from Sector Santa Rosa, ACG, Costa Rica. Folia Primatologica; International Journal of Primatology, 84(2), 102–114.

44. Perdomo, G., & Henry Dong, H. (2009). Apolipoprotein D in lipid metabolism and its functional implication in atherosclerosis and aging. Aging, 1(1), 17–27.

45. Pinto, S. L., Henriquez, M. C., Cheves Hernandez, S., Duytschaever, G., Wit, J., Avramenko, R. W., Gilleard, J. S., Orkin, J. D., & Melin, A. D. (2023). Promise and limitations of 18S genetic screening of extracted fecal DNA from wild capuchins. Frontiers in Ecology and Evolution, 11. 10.3389/fevo.2023.1176681

46. Rühlemann, M. C., Bang, C., Gogarten, J. F., Hermes, B. M., Groussin, M., Waschina, S., Poyet, M., Ulrich, M., Akoua-Koffi, C., Deschner, T., Muyembe-Tamfum, J. J., Robbins, M. M., Surbeck, M., Wittig, R. M., Zuberbühler, K., Baines, J. F., Leendertz, F. H., & Franke, A. (2024). Functional host-specific adaptation of the intestinal microbiome in hominids. Nature Communications, 15(1), 326.

47. Sadoughi, B., Hernández-Rojas, R., Hamou, H., Lopez, R., Mah, M., Slikas, E., Simmons, S., Orkin, J. D., Higham, J. P., Brosnan, S. F., Jack, K. M., Campos, F. A., Snyder-Mackler, N., & Melin, A. D. (2025). Non-invasive measures of DNA methylation capture molecular aging in wild capuchin monkeys. In bioRxivorg. 10.1101/2025.06.03.657357

48. Srivathsan, A., Ang, A., Vogler, A. P., & Meier, R. (2016). Fecal metagenomics for the simultaneous assessment of diet, parasites, and population genetics of an understudied primate. Frontiers in Zoology, 13, 17.

49. Swindell, W. R., Johnston, A., Xing, X., Little, A., Robichaud, P., Voorhees, J. J., Fisher, G., & Gudjonsson, J. E. (2013). Robust shifts in S100a9 expression with aging: a novel mechanism for chronic inflammation. Scientific Reports, 3(1), 1215.

50. Tacutu, R., Thornton, D., Johnson, E., Budovsky, A., Barardo, D., Craig, T., Diana, E., Lehmann, G., Toren, D., Wang, J., Fraifeld, V. E., & de Magalhães, J. P. (2018). Human Ageing Genomic Resources: new and updated databases. Nucleic Acids Research, 46(D1), D1083–D1090.

51. Tsutaya, T., Mackie, M., Sawafuji, R., Miyabe-Nishiwaki, T., Olsen, J. V., & Cappellini, E. (2021). Faecal proteomics as a novel method to study mammalian behaviour and physiology. Molecular Ecology Resources. 10.1111/1755-0998.13380

52. Wang, S., Song, R., Wang, Z., Jing, Z., Wang, S., & Ma, J. (2018). S100A8/A9 in inflammation. Frontiers in Immunology, 9, 1298.

53. Webb, S. E., Orkin, J. D., Williamson, R. E., & Melin, A. D. (2023). Activity budget and gut microbiota stability and flexibility across reproductive states in wild capuchin monkeys in a seasonal tropical dry forest. Animal Microbiome, 5(1), 63.

54. Williamson, R. E., Webb, S. E., Dubreuil, C., Lopez, R., Cheves Hernandez, S., Fedigan, L. M., & Melin, A. D. (2021). Sharing spaces: niche differentiation in diet and substrate use among wild capuchin monkeys. Animal Behaviour, 179, 317–338.

55. Wong, J. H., Dukes, J., Levy, R. E., Sos, B., Mason, S. E., Fong, T. S., & Weiss, E. J. (2008). Sex differences in thrombosis in mice are mediated by sex-specific growth hormone secretion patterns. The Journal of Clinical Investigation, 118(8), 2969–2978.

56. Worsley, S. F., Videvall, E., Harrison, X. A., Björk, J. R., Mazel, F., & Wanelik, K. M. (2024). Probing the functional significance of wild animal microbiomes using omics data. Functional Ecology. 10.1111/1365-2435.14650

